# Designing anti-predator training to maximize learning and efficacy assessments

**DOI:** 10.1101/2021.11.30.470590

**Authors:** Alison L. Greggor, Bryce M. Masuda, Anne C. Sabol, Ronald R. Swaisgood

## Abstract

Despite the growing need to use conservation breeding and translocations in species’ recovery, many attempts to reintroduce animals to the wild fail due to predation post-release. Released animals often lack appropriate behaviours for survival, including anti-predator responses. Anti-predator training—a method for encouraging animals to exhibit wariness and defensive responses to predators—has been used to help address this challenge with varying degrees of success. The efficacy of anti-predator training hinges on animals learning to recognize and respond to predators, but learning is rarely assessed, or interventions miss key experimental controls to document learning. An accurate measure of learning serves as a diagnostic tool for improving training if it otherwise fails to reduce predation. Here we present an experimental framework for designing anti-predator training that incorporates suitable controls to infer predator-specific learning and illustrate their use with the critically endangered Hawaiian crow, ‘alalā (*Corvus hawaiiensis*). We conducted anti-predator training within a conservation breeding facility to increase anti-predator behaviour towards a natural predator, the Hawaiian hawk, ‘io (*Buteo solitaries*). In addition to running live-predator training trials, we included two control groups, aimed at determining if responses could otherwise be due to accumulated stress and agitation, or to generalized increases in fear of movement. We found that without these control groups we may have wrongly concluded that predator-specific learning occurred. Additionally, despite generations in human care that can erode anti-predator responses, ‘alalā showed unexpectedly high levels of predatory wariness during baseline assessments. We discuss the implications of a learning-focused approach to training for managing endangered species that require improved behavioural competence for dealing with predatory threats, and the importance of understanding learning mechanisms in diagnosing behavioural problems.

## Introduction

Many conservation translocations—i.e. human-mediated relocations of wildlife to improve species’ and habitat recovery—fail despite large commitments of resources (Hoffmann et al., 2010; Seddon et al., 2014).. Many translocation failures can be attributed to predation after release (Fischer & Lindenmayer, 2000; Moseby et al., 2011), yet the behavioural mechanisms leading to increased predation are infrequently acknowledged or addressed (Berger-Tal et al., 2020). Therefore conservation interventions that reduce behavioural vulnerability to predation have the potential to improve translocation outcomes widely (Berger-Tal et al., 2020). Deficiencies in released animals’ anti-predator responses(Berger-Tal et al., 2020; Shier, 2016), are a likely contributor to post-release predation and subsequent translocation failure, especially when source populations have been free from predation pressure (Ross et al., 2019).. Just as other natural behaviours often erode in human care (Kraaijeveld-Smit et al., 2006; McPhee & Carlstead, 2010), predator-free environments foster prey naivety, which results in ineffective anti-predator behaviour (Cox & Lima 2006).

Anti-predator training—in which animals living in predator-free environments are provided opportunities to learn about predators—can be a useful tool to combat prey naivety across taxonomic groups (Griffin et al., 2000; Moehrenschlager & Lloyd, 2016; Shier & Owings, 2006; Teixeira & Young, 2014), but its efficacy in translocation contexts often goes untested (Greggor et al., 2019; Ross et al., 2019; Rowell, 2020). Accordingly, despite some successes (e.g., (Shier & Owings, 2006; van Heezik et al., 1999), training has often failed to adequately change anti-predator behaviour (Campbell & Snowdon, 2009; Jolly et al., 2020) or improve survival post-release (Moseby et al., 2012). Without being able to pinpoint where and why anti-predator training goes wrong, we lose the ability to address naivety and vulnerability to predators for translocated animals.

Anti-predator training requires manipulating animal learning. Many species naturally learn about predators during development, a process that is facilitated by experiencing a predatory cue (e.g., the sight, smell, or sound of a predator) alongside a conspecific signal of danger (e.g., an alarm call, scent, or evidence of attack) (Griffin et al., 2000). Some interventions expose animals to low levels of true predation to accurately replicate these cues and facilitate learning (e.g., (Moseby et al., 2016; Ross et al., 2019). However, losing animals to predators pre-release is often not practical when working with endangered species, with few release candidates, or due to welfare concerns. Therefore Training efforts often try to mimic the natural learning process, by pairing an aversive stimulus (e.g., a conspecific alarm cue or physical restraint) with a predator or replica (Shier & Owings, 2006; Teixeira & Young, 2014). Ideally, these presentations utilize classical conditioning learning mechanisms, allowing animals to rapidly remember the predator, not simply the context where they encountered it (Griffin et al., 2000). When successful, pre-release training enables animals to distinguish predatory threats from non-threatening stimuli in the environment and respond to actual predators in the wild, despite never having been directly attacked. Such learning is necessary for training to be effective, but measuring anti-predator learning over the course of training is not always straightforward.

A common framework for demonstrating learning compares behaviour before and after training between a trained and a control, un-trained group (Griffin et al., 2000). While this setup can successfully document changes to anti-predator behaviour over the course of training, it does not expose the root causes of behavioural change. Specifically, two cognitive mechanisms—sensitization and generalization—can falsely present as predator learning in trained groups if not adequately addressed with experimental controls, each of which may have different downstream effects for post-release survival. Sensitization occurs when animals become more responsive to repeated presentations of stimuli, and is especially likely after a mildly aversive stimulus (Shettleworth, 2010). If animals cue into aspects of the training setup and anticipate danger, sensitization during repeatedly fear-inducing training sessions could drive apparent anti-predator behaviour during training, without target animals actually learning about the predator (e.g., (Mathis & Smith, 1993). Accordingly, sensitized animals would be unlikely to engage in anti-predator behaviour when encountering a predator outside of the training setup. Second, animals may not learn about the predator itself, but simply learn a generalized fear of animacy or animate stimuli in certain situations. While responding fearfully to a broad category of animate stimuli (including towards non-predators) may help with initial survival since predators would be avoided, animals can incur energy and resource costs if they consistently over-respond to false predatory threats (Carthey & Banks, 2014). In both cases, animals that show a heightened response after training may not express optimal anti-predator responses post-release. Therefore, by including experimental controls that offer repeated presentations of fearful (but not predatory) stimuli, and controls with non-fearful, animate stimuli, training designs can document true anti-predator learning, while ruling out sensitization and generalization. These additional controls help diagnose why apparently trained animals may not experience the survival or fitness benefits training is expected to provide. Additionally, it can give managers and researchers an opportunity to assess the efficacy of training prior to release, which could prevent unnecessary deaths if training methods can be adjusted to facilitate predator-specific learning that reduces vulnerability to predators post-release.

To illustrate the importance of these cognitive considerations, we tested the efficacy of anti-predator training with the critically endangered ‘alalā, or Hawaiian crow (*Corvus hawaiiensis*). ‘Alalā went extinct in the wild in 2002 and have been the subject of intensive reintroduction efforts since 2016. Previous attempts to re-establish the species faced many challenges, including predation by their natural predator, the ‘io (Hawaiian hawk, *Buteo solitaries*) (U.S. Fish and Wildlife Service, 2009). Therefore future planning efforts incorporated the use anti-predator to improve the chances of survival (VanderWerf et al., 2013). We examined the efficacy of training in breeding facilities with ‘alalā that were not designated for imminent release, allowing us to evaluate methods with a larger, more robust sample. We measured the anti-predator responses of ‘alalā towards a predator model before and after a classical fear conditioning training, across birds that received one of three treatments: a live predator, a control fear stimulus (net), and a control animate object (live chicken). These learning-focused control treatments were designed to help identify sensitization (fear-inducing, non-predatory net) or generalization (non-fearful, animate chicken) to identify if factors other than anti-predator learning contribute to increases in anti-predator behaviour.

Our experiment was designed to differentiate among these alternative learning processes that all otherwise produce heightened anti-predator behaviour in the context of training. If the live predator group responded with greater anti-predator responses to the model in the evaluation trial than the other two groups, there would be strong evidence that the training produced predator-specific learning, and therefore may reduce the probability of predation if these birds were released. In contrast, if ‘alalā showed little increase in anti-predator behaviour in the live predator group, or showed increases across any other group, the question of training effectiveness would be more complex. If ‘alalā responded with greater anti-predator behaviour in all three groups, we would infer that training caused the birds to become sensitized to threatening/novel stimuli, instead of producing predator-specific learning. With this result we would predict that training offers little advantage in helping animals avoid predators since they did not learn to fear the predator itself. Meanwhile, if ‘alalā showed increases in anti-predator behaviour in the live predator and chicken groups only, we would conclude they acquired a learned fear response to general animacy. From a conservation management standpoint, this last outcome, which would not show predator-specific learning, could still be beneficial, depending on the energetic or resource costs ‘alalā may incur by responding unnecessarily to harmless avian or other animate stimuli. However, there are clear advantages to developing anti-predator training programs that facilitate learning to fear and display appropriate antipredator behaviour only to the predators, and not to other animate or inanimate objects that present no risk.

## Materials and Methods

### Study species

‘Alalā are the only remaining corvid species native to the Hawaiian Islands. They are a generalist forager, with one surviving native predator, the ‘io. Populations of ‘alalā declined precipitously during the late 20^th^ century due to habitat degradation, disease, human conflict and predation by non-native mammals (U.S. Fish and Wildlife Service, 2009). Despite supplementation translocations in the 1990’s, ‘alalā went extinct in the wild in 2002. Since then, conservation breeding has increased the population from fewer than 20, to over 120 in human care today. Reintroduction translocations were initiated in 2016 and are currently ongoing. However, translocation efforts have faced similar challenges over time. Predation by ‘io has been a primary cause of post-release losses in historical (U.S. Fish and Wildlife Service, 2009) and ongoing translocations (Greggor et al., 2021). Therefore the development of anti-predator training to reduce mortality from ‘io figures prominently in recovery planning documents (VanderWerf et al., 2013). Despite the need for effective training, the small size of release cohorts and critically endangered status of the species meant that training for the release birds could not contain untrained control groups with later measures of survival, in case it resulted in the deaths of untrained birds (Greggor et al., 2021). Therefore, the need for further research arose within the conservation breeding flock, where control groups would not be released and thus face no adverse survival impacts, to improve training methods in future recovery efforts.

### Birds and housing

We tested ‘alalā housed in an ongoing conservation breeding program at the Keauhou Bird Conservation Center (KBCC) in Volcano, Hawai’i. Our sample comprised hand-reared (N=35) and partially or fully parent-reared birds (N=8). We tested ‘alalā in their home enclosures, with their mate, family group, or juvenile social group. Since ‘alalā are a social species, testing individuals alone can cause stress and compromise the reliability of anti-predator responses. Moreover, training is more effective in natural social groups (Shier & Owings, 2007). Therefore, data was taken on each individual within their group and the potential for social effects was accounted for statistically with mixed effects models (see below). No ‘alalā tested in this study had previously been trained to fear ‘io as part of release efforts, but all had the occasional opportunity to observe wild ‘io that are resident in the area. It is possible they may have observed predation events by ‘io on other forest birds, but these events were never observed by staff in the years leading up to this study.

All birds had access to food and water, and were exposed to ambient light and weather conditions. Each aviary contained areas of covered perching, an indoor feeding chamber, and open areas. See (Greggor et al., 2018) for a detailed description of husbandry, enrichment and housing practices. While the dimensions of each aviary differed slightly, the basic setup was the same (Fig. 1).

**Figure 1.**
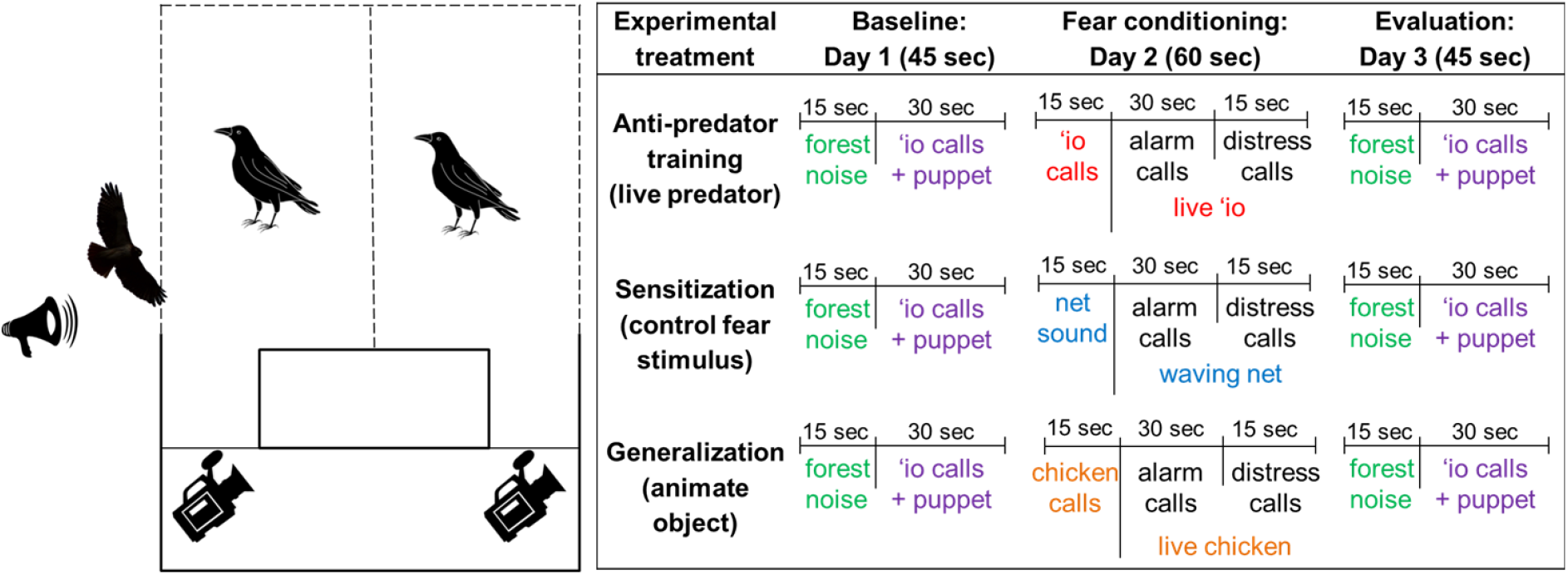
Experimental setup and schedule. In 13 of the 19 multi-chambered aviaries, individuals could move freely between the two large chambers. The speaker (~2m from aviary wall) and fear stimulus (‘io pictured) were placed on the outside of the aviary. The behaviour of birds was video recorded and observed live from inside the aviary’s observation corridor. The dotted lines indicate areas of mesh that offer visual access; areas without dotted lines are solid walls through which the birds could not see. The experimenter presented the stimuli while standing out of sight, up against the solid wall. During each trial, three minutes of behavioural data was also taken immediately prior to and after the stimuli presentation listed above. The fear conditioning day did not contain forest noise to prevent ‘alalā from associating the forest noise with the ensuing alarm and distress calls.

### Stimuli and trials

We conducted all trials from July 7-20, 2018, between 08:00-16:00. We assigned each aviary to one of three experimental treatments, exposing each bird group to a series of three trials. The three trials (baseline, fear conditioning, and evaluation) occurred on consecutive days at the same time of day. Each trial, regardless of treatment, consisted of three periods: 1) a three-minute pre-trial observation, 2) a period of stimuli exposure (whose duration and stimuli depended on the treatment and trial day), and 3) a three-minute post-trial observation (Fig. 1). Including these pre- and post-trial periods for each trial allowed us to eliminate the potential that we accidentally induced fear while setting up the trial, prior to the presentation of stimuli, which would muddy the results. The experimenter set up the trials by putting out the speaker and camcorders, allowing the birds to settle for ten minutes, and then beginning the pre-trial observation period. The experimenter then presented experimental stimuli, which lasted either 45 seconds or one minute, depending on the trial type. Once the stimuli were removed, the three-minute post-stimuli period began. All trials followed this same schedule, but the stimuli differed by trial type and treatment group (Fig. 1).

We exposed each experimental group to the same stimuli for baseline (day 1) and evaluation (day 3) trials: an audio recording and a flapping model ‘io. These recordings contained 15 seconds of ambient forest noise (recorded at KBCC)—reducing the likelihood ‘alalā would be startled by a sudden sound from the speaker—followed by 30 seconds of ‘io territory calls. We created two exemplar audio recordings to address pseudoreplication, each with a similar number and timing of calls. Subjects randomly received one exemplar for the baseline and the other for the evaluation. During trials, the hidden experimenter presented the model ‘io next to the aviary as the recording started playing ‘io calls. The model was an ‘io taxidermy, mounted on a pole with wings extended, containing a mechanism that tipped the body forward on command. The experimenter moved the model similarly for each presentation, making the dipping motion every 2 seconds and holding for 2 seconds.

During fear conditioning trials (day 2) the experimenter presented one of three stimuli, depending on the experimental treatment: a live ‘io predator (anti-predator training), a net (sensitization control), or a live chicken (generalization control). The experimenter played sounds specific to the fear stimulus for 15 seconds, and then presented the stimulus alongside an audio track containing ‘alalā alarm and distress calls. We previously recorded ‘alalā alarm calls during husbandry-related disturbance (e.g., nest-checks) and distress calls during routine veterinary procedures. The alarm calls were overlaid on each other to mimic a flock of birds (increasing the perceived risk of danger, Coomes et al. 2019), followed by a single individual emitting a series of distress calls. Like other passerines, ‘alalā make distress calls when faced with imminent danger, such as during physical capture and restraint, potentially offering information about threats to others (Griffin, 2008). No calls from experimental subjects in this study were used in the audio files.

For the live ‘io treatment we borrowed a glove-trained ‘io from the Panaewa Rainforest Zoo and Gardens in Hilo, Hawai’i. He was maintained at the KBCC in an outdoor enclosure between trials. He voluntarily stepped onto a falconer’s glove for each presentation and was encouraged to flap at the ‘alalā, in response to gentle motion of the glove. The handler remained hidden behind the side of the aviary, extending the glove into the area of visual access for the ‘alalā. The audio track used prior to the live ‘io presentation was a separate compilation of ‘io territorial calls than those used for baseline or evaluation trials.

For the sensitization control treatment, we presented a large black recapture net during the fear conditioning stimuli. As the primary recapture method at the facility, the net served as an artificial, fear-inducing stimulus. Before presenting the net, the experimenter clanked two net poles together repeatedly (every 2 seconds) for 15 seconds, simulating the sound of staff removing nets from work trucks during recapture. Immediately after the net sounds, the hidden experimenter played the ‘alalā alarm and distress calls, while waving the net in a similar range of movement to the ‘io’s flapping wings.

For the generalized animacy control treatment, we presented a live chicken during the fear conditioning trial. The ‘alalā had never seen a live adult chicken, and have no evolutionary history of predation by ground-based birds, so we predicted that it would not elicit anti-predator responses (although its novelty could still elicit neophobia, e.g., (Greggor et al., 2020). Prior to the presentation of the chicken, the experimenter played a series of non-fear related chicken calls. The hidden experimenter then played the ‘alalā alarm calls and held the chicken out on the side of the aviary, encouraging him to flap.

We edited sound files using Audacity® software and broadcast them from an Altec Lansing Bluetooth speaker. We broadcast sound files at the same maximum volume level (80 dB), verified with the Decibel X Power Meter app for iPhones. Unless otherwise specified, we collected sound recordings using a Roland R-05 acoustic recorder and a Sennheiser microphone with a Rycote softie wind cover. Data collection and analysis

We collected behavioural data via live observer and video recorded from multiple angles. We recorded the number of anti-predator behaviours (alarm calls and rapid escape flights across the length of the aviary) and non-fear behaviours (affiliative begging calls) across each trial period (pre-stimuli, during stimuli, post-stimuli). Additionally, we classified birds’ level of engagement with the stimuli into one of several categories (Table 1). An independent observer, blind to the experimental questions and original data, recoded a subset of videos (20%) to assess inter-observer reliability, which was calculated with an intraclass correlation coefficient (ICC).

**Table 1.**
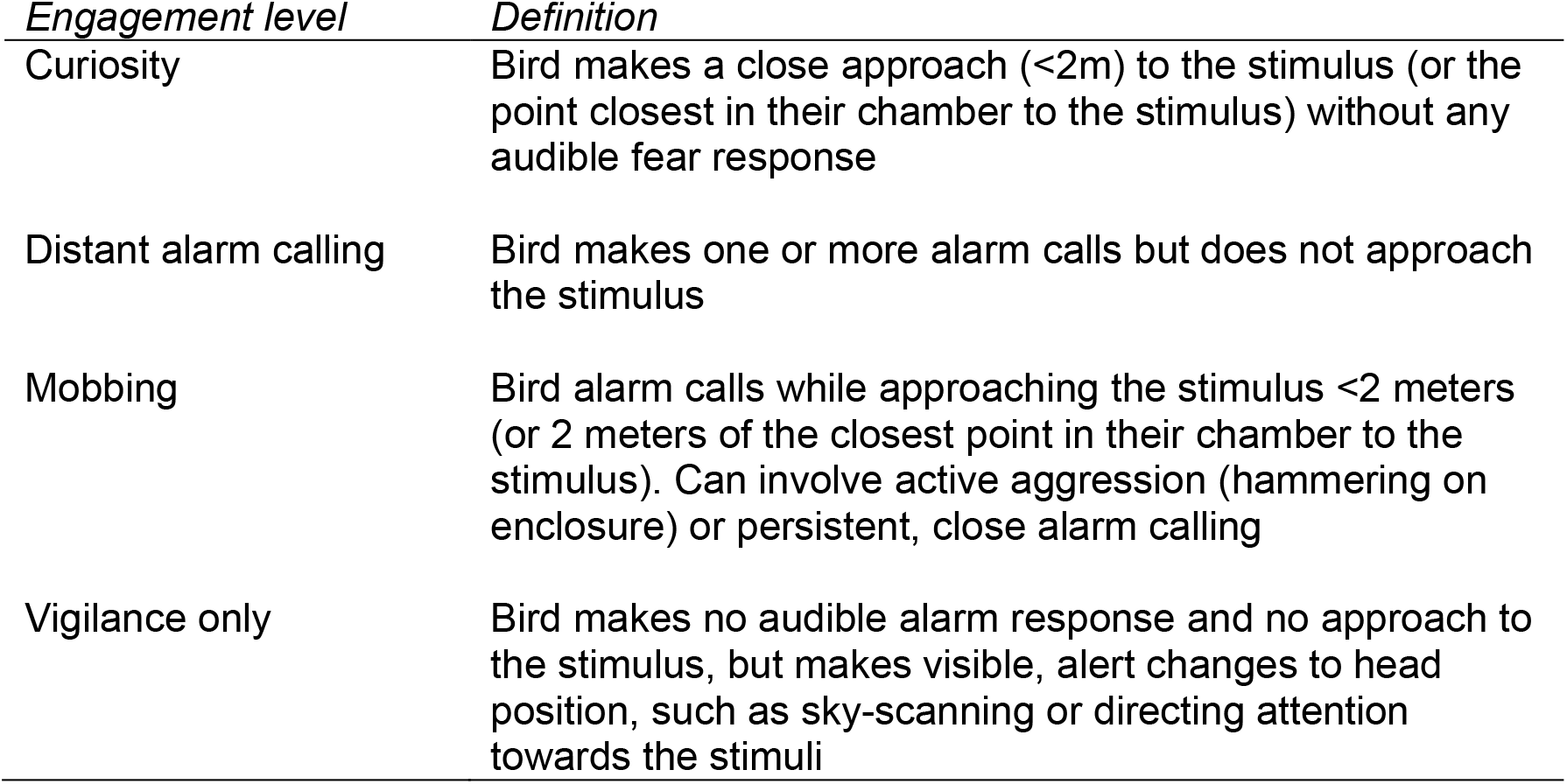
Definitions of behavioural categories used to explain levels of approach behaviour during exposure to the experimental stimuli.

| *Engagement level* | *Definition* |
| --- | --- |
| Curiosity | Bird makes a close approach (<2m) to the stimulus (or the point closest in their chamber to the stimulus) without any audible fear response |
| Distant alarm calling | Bird makes one or more alarm calls but does not approach the stimulus |
| Mobbing | Bird alarm calls while approaching the stimulus <2 meters (or 2 meters of the closest point in their chamber to the stimulus). Can involve active aggression (hammering on enclosure) or persistent, close alarm calling |
| Vigilance only | Bird makes no audible alarm response and no approach to the stimulus, but makes visible, alert changes to head position, such as sky-scanning or directing attention towards the stimuli |

Non-fear behaviour occurred too infrequently (1.4% of trial periods) to merit formal analysis. Therefore, we focused on anti-predator behaviour and level of engagement with the stimuli to answer three main questions. First, to confirm that all treatment groups began with similar levels of fear towards the ‘io model, we compared anti-predator behaviours across treatments on the baseline trial (day 1). Second, to determine if the live predator, net, and chicken stimuli elicited different responses during the fear-conditioning trial, we compared anti-predator behaviour and engagement levels across treatments during fear conditioning (day 2). Finally, we compared changes between baseline trials (day 1) and evaluation trials (day 3) across treatments to determine if anti-predator behaviour increased over trials to indicate predator-specific learning.

We compared anti-predator behaviour for each question with a generalized linear mixed model (GLMM) in R version 3.6.2 (Team, 2019) with the lme4 package (Bates et al., 2013). Each model contained the following explanatory variables: sex, number of birds present in the aviary, whether the bird had access to the stimuli side of the aviary, trial period (pre-stimuli, stimulus, post-stimuli), and treatment (anti-predator training, sensitization control and generalization control). Models examining changes to anti-predator behaviour over time also contained an interaction between trial number (baseline = 1, evaluation = 3) and treatment. Subject ID was included as a random factor for all models. Accounting for this source of non-independence was necessary since each individual contributed data points for the three trial periods, across all three trials. Age was not included since only one aviary contained birds younger than breeding age, and their data were not outliers. Parent-versus hand-reared birds were approximately evenly spread across treatments, and later inclusion of rearing type did not improve any final models, so this distinction was not included in main analyses (Table S4). The relative influence of factors was assessed with AICc values (calculated with the MuMIn package, (Barton, 2020) due to the relatively small sample size, and factors were dropped from the model if their inclusion did not reduce AICc values by >2. Model fit and assumptions, including dispersion, outliers, uniformity and zero inflation were checked with the DHARMa package (Hartig, 2021), and data transformations and model error structure chosen accordingly.

We transformed anti-predator behaviours to a binomial variable (presence of anti-predator behaviour = 1, absence = 0) for comparing baseline responses across treatments. In analysing behaviour across treatments during the fear conditioning trial, we converted behavioural sums (alarm calls plus full flights) to a count per three-minute period, to account for the different trial period durations, and analysed them with a negative binomial error distribution. For the third question evaluating changes in anti-predator behaviour from the baseline to evaluation trials, we focused on the “during” stimuli period. We converted anti-predator behaviour to a count per three-minute period, square-root transformed it and analysed it with a Gaussian error distribution. We conducted a post-hoc power analysis on the GLMM for the during-stimuli comparison of baseline and evaluation trials using the simr package (Green & Macleod, 2016).

We conducted separate analyses to investigate birds’ engagement levels with the fear stimuli. We examined whether birds were more likely to respond within a certain category (Table 1) in the fear-conditioning trial (day 2) with an exact multinomial test, assuming an equal 25% chance of any behaviour occurring. We ran post hoc binomial tests with Bonferroni corrections to investigate categories of interest. We also compared birds’ response between baseline and evaluation trials with a marginal homogeneity test, using the coin package (Zeileis et al., 2008).

### Ethical statement

This work using animal subjects was approved by San Diego Zoo Wildlife Alliance’s IACUC committee (No. 16-009). Permits allowed conservation breeding of ‘alalā (USFWS Native Endangered Species Recovery Permit TE060179-5, State of Hawaii Protected Wildlife Permit WL19-16) and the possession of live ‘io (USFWS Special Purpose Miscellaneous permit MB09204C-1).

## Results

We tested 43 ‘alalā (N=13 chicken, 13 live predator, 17 net) across 19 aviaries. Of these, two aviaries were excluded for fear conditioning and evaluation trials (days 2 and 3) because birds in one aviary started breeding (net treatment), and in another the speaker malfunctioned midway through the alarm and distress calls (chicken treatment). Inter-observer reliability was high for the composite measure of alarm calls and full flights (ICC(1)=0.91, p<0.001, CI=0.86-0.94), and for the level of engagement (96.4% concurrence). One bird was not reliably visible on video during the fear conditioning trial (day 2), and her data was removed for that day.

### Treatment effect during baseline trials

In baseline trials (day 1), all treatment groups showed increased anti-predator during the ‘io model presentation and post-stimuli time periods in comparison to pre-stimuli anti-predator rates (Binomial GLMM, N=129 observations, 43 birds across 3 periods; During: *B*=2.35±0.67, z=3.49; Post-stimuli: *B*=1.46±0.62, z=2.34, ΔAICc= −10.428 relative to final model excluding this term; Fig. 2). Birds tested in larger social groups had slightly higher levels of anti-predator responses (*B*=0.87±0.43, z=2.02, ΔAICc= −2.21). There was no effect of treatment, as expected, since all groups were exposed to the same stimuli. Also, we found no effect of sex or bird access to the stimuli side of the aviary (Table S1).

**Figure 2.**
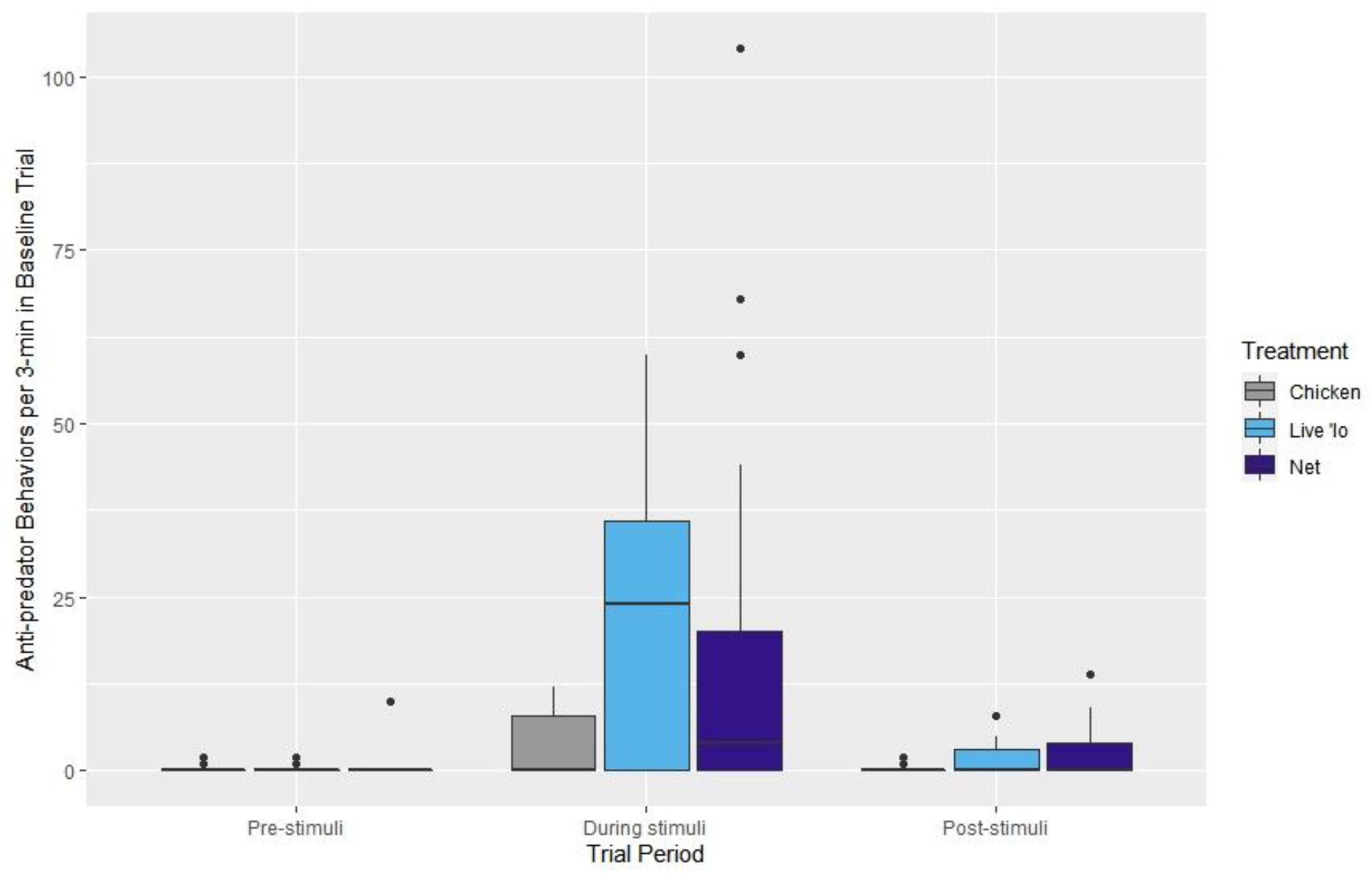
Boxplots depicting raw rates of individual anti-predator behaviour across trial periods of baseline trials (day 1). Outlying data points are pictured and horizontal line in each box depicts the median value. Anti-predator behaviour is measured as the rate of alarm calls and pace flies per minute of trial period. All treatments received the same stimuli during these baseline trials: a three-minute pre-stimuli period, 45 seconds of exposure to an ‘io model and ‘io calls, and a three-minute post-stimuli period. There was no effect of experimental treatment and all conditions show the same pattern: little anti-predator behaviour during the pre-trial, increased fear while the model and calls were present, and reduced fear once stimuli were removed in the post-trial period.

### Treatment effect during fear-conditioning trials

Similar to baseline trials, during the fear conditioning trials (day 2) ‘alalā displayed higher rates of anti-predator behaviour during the stimuli presentation and post-stimuli period (GLMM, N=111 observations, 37 birds across 3 periods, During: *B*=3.82±0.16, z=23.98; Post-stimuli: *B*=2.13±0.16, z=12.79, ΔAICc=-2737.30) relative to the pre-stimuli period. All treatment groups were indistinguishable in their rates of anti-predator behaviour, despite receiving either a live predator, net, or live chicken. However, birds in larger social groups had slightly higher rates of anti-predator behaviour (*B*=0.69±0.31, z=2.55, ΔAICc=-2.61). There was no effect of sex (Table S2). Additionally, while stimuli were present, ‘alalā were more likely to mob (Binomial test, Bonferroni correction applied, p<0.001) and less likely to exhibit curiosity (p<0.001; Figure 3).

**Figure 3.**
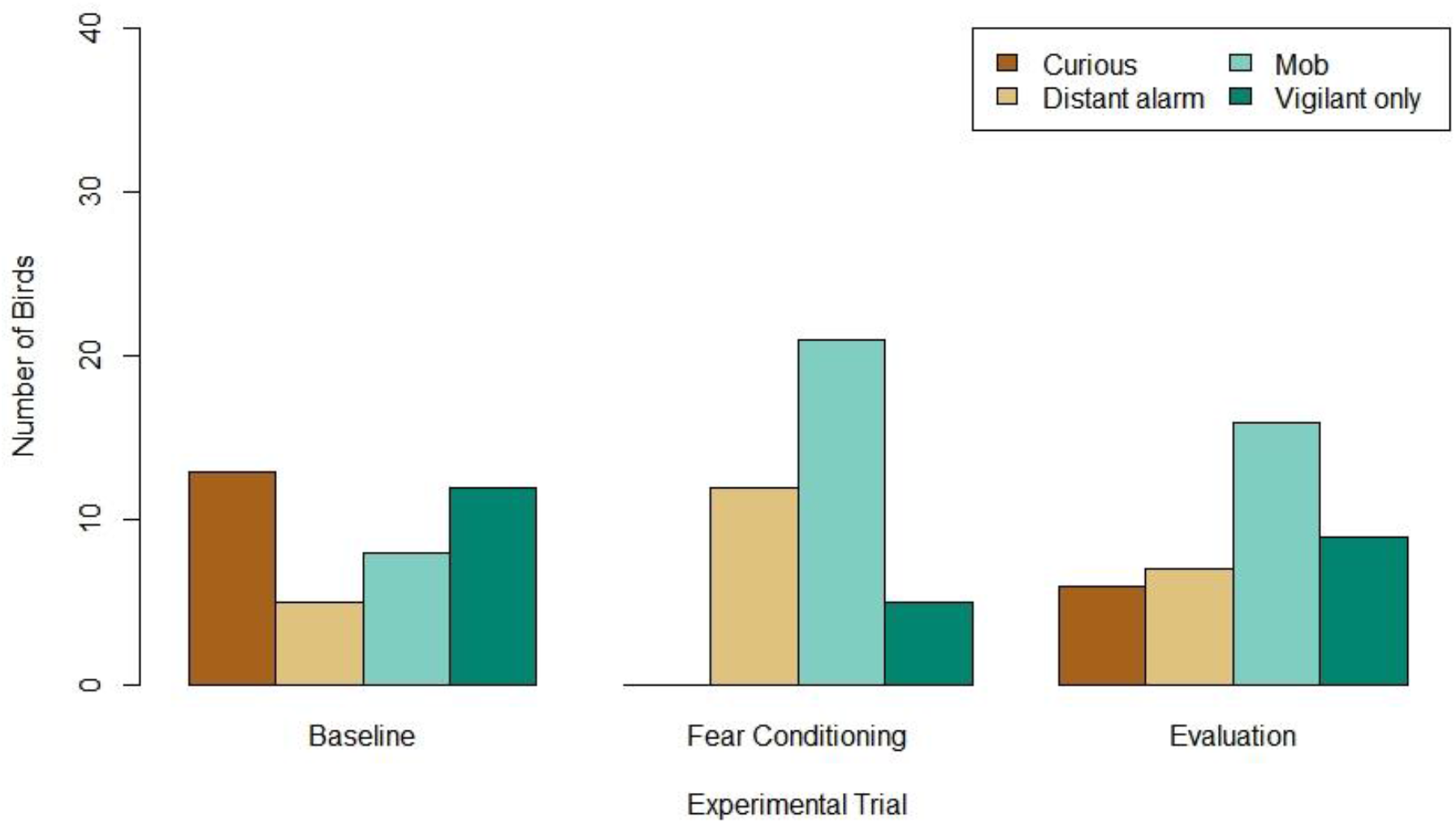
Number of ‘alalā exhibiting response categories to the stimuli during each trial type. During the Fear Conditioning trials, birds were more likely than chance to mob and less likely than chance to show a curious response. In comparing between trial types, more ‘alalā mobbed the taxidermy ‘io model during the Evaluation than Baseline trials.

### Treatment effect on predator-specific learning (baseline versus evaluation)

There was no interaction between treatment and trial (Table 2). Across all treatments, ‘alalā increased their anti-predator behaviour towards the ‘io model during the evaluation trials, relative to baseline trials (*B*=1.51±0.58, z=2.62, ΔAICc=-4.87; Figure 4). ‘Alalā tested in larger group sizes showed higher levels of anti-predator behaviour (*B*=1.95±0.62, z=3.13, ΔAICc=-7.48). Additionally, more ‘alalā mobbed the ‘io model during evaluation than baseline trials (Marginal Homogeneity test; χ^2^=8.027, df=3, p=0.045; Figure 3). Power analyses revealed a low likelihood of finding an effect during the stimuli presentations, even for relatively large effect sizes, (*B*=4.0, power=62.90%, CI=59.82-65.90%; *B*=2.0, power=23.20%, CI=20.62-25.94%; *B*=1.0, power=7.20% CI=5.68-8.98%). However, the data illustrate similar upward trends from baseline to evaluation in each treatment group (Fig. 4), showing little support for an interaction between treatment and trial, which would have indicated predator-specific learning.

**Table 2.** Results of model selection process for GLMM on anti-predator behaviour during the presentation of the ‘io model with data from the baseline and evaluation Trial (day 1 and 3). The experimental Treatment denoted groups that experienced either a live ‘io, net or live chicken. The model term Access denoted whether birds were housed in the chamber adjacent to the experimental stimuli. ^a^The change in AICc represents the difference in AICc value from the model listed directly above. Terms were included in the model if dropping them increased AICc by more than 2. ^b^Final model.

| *Model* | *$\Delta AICc^a$* |
| --- | --- |
| Anti-predator behaviour ~ Num_birds + Trial + Treatment + Access + Sex Treatment + Treatment: Trial + (1\|Bird) | 0.0 |
| Anti-predator behaviour ~ Num_birds + Trial + Treatment + Access + Sex Treatment + (1\|Bird) | -2.49 |
| Anti-predator behaviour ~ Num_birds + Trial + Treatment + Access + (1\|Bird) | -1.38 |
| <sup>b</sup>Anti-predator behaviour ~ Num_birds + Trial + Treatment + (1\|Bird) | -2.11 |
| Anti-predator behaviour ~ Trial + (1\|Bird) | +6.10 |
| Anti-predator behaviour ~ Num_birds + (1\|Bird) | +3.48 |

**Figure 4.**
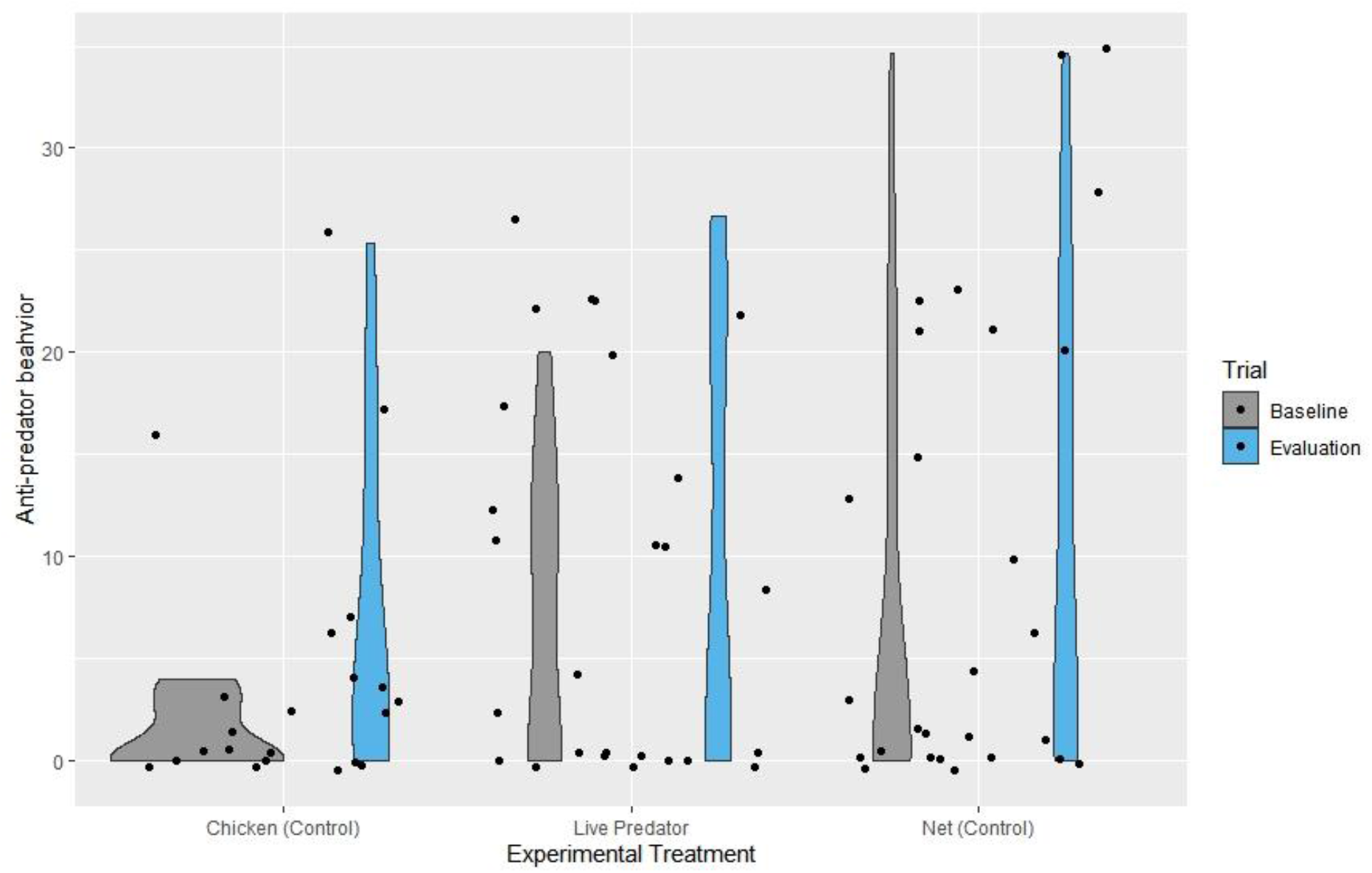
Violin plots and jittered raw datapoints depicting anti-predator behaviour on the baseline (day 1) and evaluation (day 3) trials across experimental treatments, while stimuli were present. Anti-predator behaviour is depicted here as the number of alarm calls and full aviary flights during the stimuli period. Birds in all experimental conditions demonstrated an increase in anti-predator behaviour during the evaluation in comparison to baseline trials, and no interaction effect was detected statistically.

## Discussion

We illustrated a learning-focused approach to testing anti-predator training, incorporating suitable controls to determine whether predator-specific learning occurred. Using this approach, we measured the efficacy of training for increasing anti-predator behaviour in ‘alalā towards their natural predator. While we documented increases in anti-predator behaviours after training, our control treatments also saw increases, suggesting that factors other than anti-predator learning were in play. Also, we found that ‘alalā already showed substantial anti-predator wariness towards ‘io prior to experiencing training, and that birds displayed more anti-predator behaviour when tested in larger groups. Our results demonstrate the difficulties in designing anti-predator training and emphasize the importance of considering alternative cognitive mechanisms underlying anti-predator behaviour.

Had we not conducted multiple learning controls, we may have wrongly concluded that predator-specific learning occurred in ‘alalā exposed to live predator training. Many published accounts of training include only a non-training control to support their efficacy (Crane & Mathis, 2011; Teixeira & Young, 2014). Yet, we have shown that several alternative cognitive hypotheses could explain a similar increase in fear after exposure to training and control stimuli. ‘Alalā displayed more anti-predator behaviour toward an ‘io model after the live predator and either control treatment, suggesting that they became sensitized to the setup and anticipated the appearance of dangerous stimuli. In other words, the initial model ‘io may have primed the birds to find the next presentation of the model scarier than before (the opposite of habituation, (Shettleworth, 2010). Alternatively, experiencing multiple days of fear stimuli may have put the birds in a sustained agitated state (e.g., (McIvor et al., 2018), causing them to react more strongly to the predator model over time. In future, work with ‘alalā and other species being trained for reintroduction could extend the time between baseline and evaluation trials or conduct the evaluation trials in a different enclosure (e.g., (Mathis & Smith, 1993), potentially reducing the likelihood that animals carry over motivational effects between trials. This would remove one potential source of sensitization, and also reduce the likelihood that other non-learning factors, such as neophobia, contribute to anti-predator responses (Abudayah & Mathis, 2016).

We faced a challenge that is common to many translocation programs; there are often few animals available for testing. Even with the relatively large sample size for studies of this nature (N=43, representing approximately 1/3 of the species), our analysis lacked sufficient power to confirm that birds responded statistically similarly to live predator and control treatments. However, we remain confident in our general findings because numerical trends suggested similar increases in anti-predator behaviour across treatments. By adding multiple controls to our setup, we increased the effort and sample size needed, but were able to better assess the efficacy of our training. Had we just included the live chicken control, we would have been unable to determine if ‘alalā generalized their responses to animate avian stimuli, or if they sensitized to the setup, and either conclusion could lead to different survival outcomes. Specifically, if birds were merely sensitized to the training, they would not likely respond with anti-predator behaviour to actual predators post-release, because the diverse contexts where they encounter predators would not mirror the exact training setup.

Even if conducting two types of controls is not possible, there are other benefits to approaching training with learning in mind. For example, anti-predator training relies on tapping into social cues and reliable predator-relevant cues (Griffin, 2004; Shier & Owings, 2007), but the social environment and types of cues used can both influence learning outcomes. The social environment contributed to the expression of anti-predator behaviour we documented, with larger groups of birds demonstrating more anti-predator behaviour in all trials. However, our findings do not indicate that group size influenced learning. Whether the training was more effective in larger ‘alalā groups requires further evaluation, but in other species matching natural social groupings improves training outcomes (Shier & Owings, 2007). Additionally, alarm calls are potent cues for corvids (Coomes et al., 2019), and the calls we played during fear conditioning trials proved highly effective in producing fear, even without predators present. For instance, the chicken was intended to be a non-threatening, animate stimulus, but was unexpectedly fear-inducing for ‘alalā. Other birds have learned to fear novel and otherwise non-threatening animals and inanimate objects when paired with alarm calls (Curio, 1978), which may have occurred in our trials.

Documenting learning requires a baseline measure of behaviour, which can also help assess training needs. ‘Alalā showed high levels of anti-predator behaviour towards the model predator during baseline assessments, corroborating and strengthening similar evidence from a separate study on a smaller set of juvenile released ‘alalā (Greggor et al., 2021). Together, these results suggest that much of the species is not naïve to ‘io as a threat, despite their generations in human care. It is unclear whether their baseline predatory wariness stems from observing the few resident ‘io near the facility, or has been maintained through multiple generations in conservation breeding (contrary to other species’ declines in anti-predator behaviour, e.g., (Kraaijeveld-Smit et al., 2006). Although vulnerability to predators may have contributed to losses seen in historical (U.S. Fish and Wildlife Service, 2009) and recent translocations (Greggor et al., 2021), ‘alalā’s high baseline anti-predator responses suggest that ‘alalā recognize ‘io as a threat. Other aspects of predatory evasion—such failing to act appropriately after recognizing a predator, vigilance levels in the absence of a detected predator, or using habitat in ways that minimize vulnerability—may present greater issues (Greggor et al., 2021). Systematically addressing these components of predation risk and other threats post-release contributes to an adaptive management approach, while also improving the theory and application of translocation biology (Seddon et al., 2007; Taylor et al., 2017). An additional advantage of documenting baseline behaviour in anti-predator training is that it can be a screening tool to evaluate individual competency, can be compared with target behaviours of wild individuals, and allows for assessing inter-individual variation.

Ideally released animals respond fearfully to predators and only to predators. We have shown the difficulty, yet necessity of approaching anti-predator training from a cognitive standpoint to document this type of predator-specific learning. Concurrently, however, more work is needed to determine if other processes, such as sensitization or generalization, can still provide benefits, post-release. Even simple sensitization could make less-reactive animals more wary and vigilant if triggered close to release, which could be beneficial, depending on the prevalence of predators. However, generalized fear responses could levy serious costs if they divert attention from other fitness-enhancing activities (Carthey & Banks, 2014). Training controls that allow managers to identify vulnerabilities to over-responding may explain unplanned losses of animals and allow release programs greater effectiveness in adaptively managing training and post-release conditions in future. Focusing on an evidence-based approach to translocation biology offers promise for improving outcomes and prioritizing interventions with higher probabilities of success, including for behavioural-based interventions such as pre-release training (Berger-Tal et al., 2020; Seddon et al., 2007). Therefore, testing the efficacy of actions involved in translocations, especially those are labour intensive, such as anti-predator training, will help support greater conservation progress overall.

## Supporting information

Data for Greggor et al_Effectiveness of anti-pred training

R code for Greggor et al_Effectiveness of anti-pred training

## Data Statement

All data and code used for the analysis in this paper is contained within the supplementary material.

## Acknowledgements

We thank the ‘Alalā Working Group for discussions about anti-predator training, and are grateful to P. Mizuno for lending us “Kapono” from Panaewa Zoo, to D. Alverson for lending us his chicken “Homie”, to K. Earnest, K.R. Bergfeld, and K. Whitaker for help with data collection and video coding, to J. Gaudiosa-Levita for helping acquire the model ‘io, to P. Hart for lending audio recording equipment and to the dedicated animal care team at KBCC for facilitating the project.

